# Abundance compensates kinetics: Similar effect of dopamine signals on D1 and D2 receptor populations

**DOI:** 10.1101/444984

**Authors:** Lars Hunger, Arvind Kumar, Robert Schmidt

## Abstract

The neuromodulator dopamine plays a key role in motivation, reward-related learning and normal motor function. The different affinity of striatal D1 and D2 dopamine receptor types has been argued to constrain the D1 and D2 signalling pathways to phasic and tonic dopamine signals, respectively. However, this view assumes that dopamine receptor kinetics are instantaneous so that the time courses of changes in dopamine concentration and changes in receptor occupation are basically identical. Here we developed a neurochemical model of dopamine receptor binding taking into account the different kinetics and abundance of D1 and D2 receptors in the striatum. Testing a large range of behaviorally-relevant dopamine signals, we found that the D1 and D2 dopamine receptor populations responded very similarly to tonic and phasic dopamine signals. Furthermore, due to slow unbinding rates, both receptor populations integrated dopamine signals over a timescale of minutes. Our model provides a description of how physiological dopamine signals translate into changes in dopamine receptor occupation in the striatum, and explains why dopamine ramps are an effective signal to occupy dopamine receptors. Overall, our model points to the importance of taking into account receptor kinetics for functional considerations of dopamine signalling.

**Significance statement:** Current models of basal ganglia function are often based on a distinction of two types of dopamine receptors, D1 and D2, with low and high affinity, respectively. Thereby, phasic dopamine signals are believed to mostly affect striatal neurons with D1 receptors, and tonic dopamine signals are believed to mostly affect striatal neurons with D2 receptors. This view does not take into account the rates for the binding and unbinding of dopamine to D1 and D2 receptors. By incorporating these kinetics into a computational model we show that D1 and D2 receptors both respond to phasic and tonic dopamine signals. This has implications for the processing of reward-related and motivational signals in the basal ganglia.

## Introduction

The neuromodulator dopamine (DA) plays a key role in motivation, reward-related learning and normal motor function. Many aspects of DA function are mediated by its complex effects on the excitability (Day et al., 2008) and strength of cortico-striatal inputs (Reynolds et al., 2001) in the context of motor control (Syed et al., 2016), action-selection (Redgrave et al., 2010), reinforcement learning (Schultz, 2007), and addiction (Everitt and Robbins, 2005). The striatal DA concentration ([DA]) can change over multiple timescales (Schultz, 2007). Fast, abrupt increases in [DA] lasting for ≈ 1 — 3s result from phasic bursts in DA neurons (Roitman et al., 2004), which signal reward-related information (Schultz, 2007; Grace et al., 2007). Slightly slower [DA] ramps occur as animals approach a goal location (Howe et al., 2013) or perform a reinforcement learning task (Hamid et al., 2016). Finally, slow tonic spontaneous firing of DA neurons may control the baseline [DA] and change on a timescale of minutes or longer (Grace et al., 2007). However, whether e.g. learning and motivation are mediated by different timescales of DA cell firing (Niv et al., 2007) has recently been challenged (Berke, 2018; Mohebi et al., 2019). The issue of DA signalling time scales is important because the two main types of DA receptors, D1 and D2, may react to different timescales of the DA signal because of their different affinities for DA.

Based on the different DA affinities of D1 and D2 receptors (D1R and D2R), it is often assumed that striatal medium spiny neurons (MSNs) respond differently to tonic and phasic DA signals, depending on which DA receptor type they predominantly express (Dreyer et al., 2010; Surmeier et al., 2007; Grace et al., 2007; Schultz, 2007; Frank and O’Reilly, 2006). According to this “affinity-based” model, the low affinity D1Rs (i.e. with a high dissociation constant 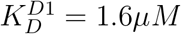; Richfield et al., 1989) cannot detect tonic changes in [DA] because the fraction of occupied D1Rs is too small (≈ 1%) at a baseline [DA] of 20*nM* and does not change much during tonic, low amplitude [DA] changes. However, D1Rs can detect phasic, high amplitude [DA] increases because they only saturate at a very high [DA]. By contrast, D2Rs have a high affinity (i.e. a low dissociation constant 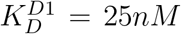; Richfield et al., 1989) leading to ≈ 40% of D2Rs being occupied at a baseline [DA] of 20nM. Due to their high affinity, D2Rs can detect low amplitude, tonic increases/decreases in [DA]. However, as D2Rs saturate at relatively low 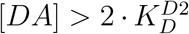, and are therefore unable to detect high amplitude, phasic increases in [DA]. This suggests that D1 and D2 type MSNs respond differently to phasic and tonic changes in [DA], solely because of the different affinities of D1Rs and D2Rs (Schultz, 2007). However, this model neglects other factors relevant for receptor occupation and is also incompatible with recent findings that D2R expressing MSNs can detect phasic changes in [DA] (Marcott et al., 2014; Yapo et al., 2017).

The affinity-based model assumes that the reaction equilibrium is reached instantaneously, whereby the receptor binding affinity can be used to approximate the fraction of receptors bound to DA. However, this assumption holds only if the receptor kinetics are fast compared to the timescale of the DA signal, which is typically not the case. For instance, D1Rs and D2Rs unbind from DA with a half-life time of *t*_1/2_ ≈ 80s (Burt et al., 1976; Sano et al., 1979; Maeno, 1982; Nishikori et al., 1980), much longer than phasic signals of a few seconds (Robinson et al., 2001; Schultz, 2007; Hamid et al., 2016). Moreover, the fraction of bound receptors might be a misleading measure for the effect of DA signals, since the abundances of D1R and D2R in the striatum are quite different. To investigate the role of receptor kinetics and abundances for DA signalling in the striatum, we developed a neurochemical model of incorporating kinetics and abundances of D1Rs and D2Rs and re-evaluated current views on DA signalling in the striatum. We show that when receptor kinetic timescales are slower than, or comparable to, the DA signalling timescales, the response of D1 and D2 DA receptor populations is similar to each other for both phasic and tonic inputs.

## Methods and Materials

### Code Accessibility

All models were implemented in Python. The models and all scripts used to generate the data and figures can be accessed here: https://bitbucket.org/Narur/abundance_kinetics/src/.

### Kinetics model

In the affinity-based model the receptor kinetics are instantaneous, so that the fraction of occupied D1 and D2 receptors (*f*_*D*1_ and *f*_*D*2_) are calculated directly from the concentration of free DA in the extracellular space, [DA], and the dissociation constant *K_D_* (see e.g. Copeland 2004):

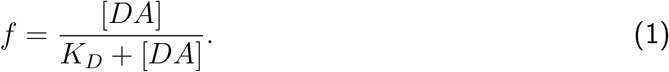

However, the dissociation constant is an equilibrium constant, so it should only be used for calculating the receptor occupancy when the duration of the DA signal is longer than the time needed to reach the equilibrium. As this is typically not the case for phasic DA signals, since the half-life time of receptors is longer (Burt et al., 1976; Sano et al., 1979; Maeno, 1982; Nishikori et al., 1980) than the timeframe of phasic signaling (Roitman et al., 2004), we developed a model which incorporates slow kinetics. When DA and one of its receptors are both present in a solution they constantly bind and unbind. During the binding process a receptor ligand complex (here called DA-D1 or DA-D2) is formed (see e.g. Copeland 2004). We refer to the receptor ligand complex as an occupied DA receptor. Below we provide the model equations for D1 receptors, but the same equations apply for D2 receptors (with different kinetic parameters). In a solution binding occurs when receptor and ligand meet due to diffusion, with high enough energy and a suitable orientation, described as:

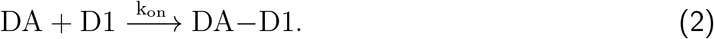

Accordingly, unbinding of the complex is denoted as:

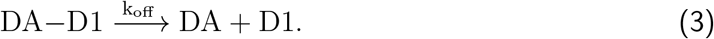

The kinetics of this binding and unbinding, treated here as first-order reactions, are governed by the rate constants *k*_on_ and *k*_off_ that are specific for a receptor ligand pair and temperature dependent. Since both processes are happening simultaneously we can write this as:

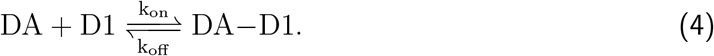

The rate at which the receptor is occupied depends on [DA], the concentration of free receptor [D1] and the binding rate constant *k*_on_:

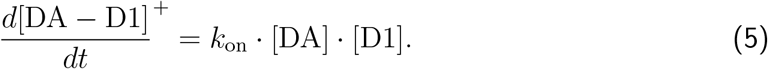

The rate at which the receptor-ligand complex unbinds is given by the concentration of the complex [DA — D1] and the unbinding rate constant *k*_off_:

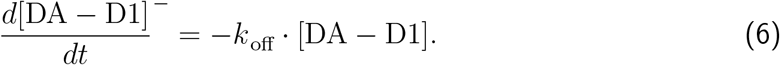

The equilibrium is reached when the binding and unbinding rates are equal, so by combining Eq. 5 and Eq. 6 we obtain:

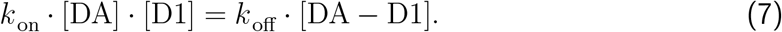

At the equilibrium the dissociation constant *K_D_* is defined as:

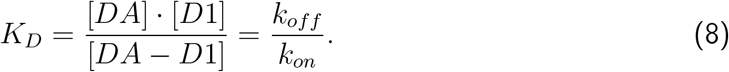

When half of the receptors are occupied, i.e. [*DA* – *D*1] = [*D*1], Eq. 8 simplifies to *K_D_* = [*DA*]. So at equilibrium, *K_D_* is the ligand concentration at which half of the receptors are occupied.

Importantly, for fast changes in [DA] (i.e. over seconds) it takes some time until the changed binding (Eq. 5) and unbinding rates (Eq. 6) are balanced, so the new equilibrium will not be reached instantly. The timescale in which equilibrium is reached can be estimated from the half-life time of the bound receptor. The half-life time assumes an exponential decay process as described in Eq. 6 and is the time required so that half of the currently bound receptors unbind. If [*DA*] = 0, and there is no more binding, the half life time of the receptors can be calculated from the off-rate by using *t*_1/2_ = *ln*(2)/*k_off_*. Signal durations should be of the same order of magnitude (or longer) than the half-life time in order for the affinity-based model with instant kinetics to be applicable.

We calculated the time course of occupied receptor after an abrupt change in [DA] by integrating the rate equation, given by the sum of Eq. 5 and Eq. 6:

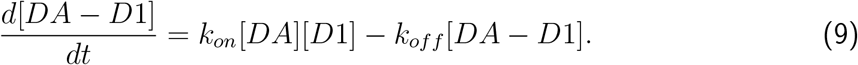

To integrate Eq. 9 we substitute

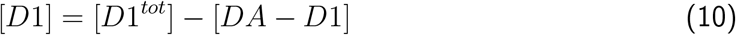

where [*D*1^*tot*^] is the total amount of D1 receptor (bound and unbound to DA) on the cell membranes available for binding to extracellular DA.

To model the effect of phasic changes in [DA] we choose the initial receptor occupancy [*DA* – *D*1](*t* = 0) = [*DA* — *D*1]^0^ and the receptor occupancy for the new equilibrium at time infinity [*DA* – *D*1](*t* = <*x*>) = [*DA* – *D*1]^∞^ as the boundary conditions. With these boundary conditions we get an analytic expression for the time evolution of the receptor occupancy under the assumption that binding to the receptor does not significantly change the free [DA]:

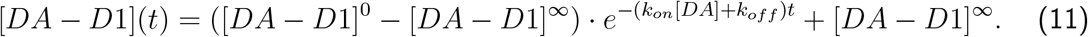

For arbitrary DA time courses we solve Eq. 9 for each receptor type numerically employing a 4th order Runge Kutta solver with a 1 ms time resolution.

We did not take into account the change in [DA] caused by the binding and unbinding to the receptors since the rates at which DA is removed from the system by binding to the receptors is much slower than the rate of DA being removed from the system by uptake through DA transporters. For example, the rate at which DA binds to the receptors is:

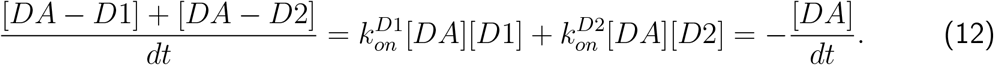

For [*DA*] = 1*μM* with a D1 and D2 occupancy of [*DA* — *D*1] ~ 20.0nM and [*DA* — *D*2] ~ 40nM (the equilibrium values for the baseline [*DA*] = 20*nM*) and 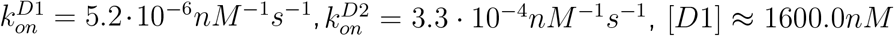 and [*D*2] ≈ 40.0*nM* the rate of DA removal through binding to the receptors is:

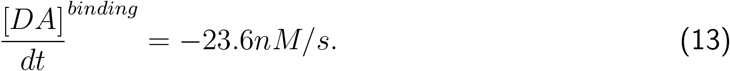

However, the DA removal rate by Michaelis-Menten uptake through the DA transporters at this concentration would be:

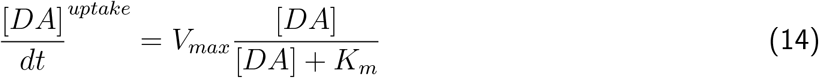

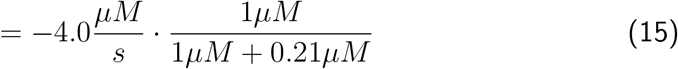

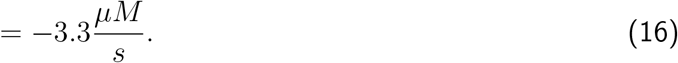

*V_max_* is the maximal uptake rate, and *K_m_* the Michaelis-Menten constant describing the [*DA*] concentration at which uptake is at half the maximum rate. As 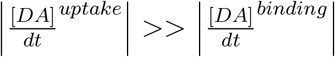, the DA dynamics are dominated by the uptake process and not by binding to the receptors. Therefore, we neglected the receptor-ligand binding for the DA dynamics in our model. However, for faster DA receptors this effect would become more important.

### Receptor parameters

An important model parameter is the total concentration of the D1 and D2 receptors on the membrane ([*D*1]^*tot*^ and [*D*2]^tot^) that can bind to DA in the extracellular space of the striatum. Our estimate of [*D*1]^tot^ and [*D*2]^tot^ is based on radioligand binding studies in the rostral striatum (Richfield et al., 1989, 1987). We use the following equation, in which *X* is a placeholder for the respective receptor type, to calculate these concentrations.

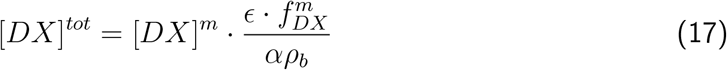

The experimental measurements provide us with the number of receptors per unit of protein weight [*D*1]^*m*^ and [*D*2]^*m*^. To transform these measurements into molar concentrations for our simulations, we multiply by the protein content of the wet weight of the rat caudate nucleus e, which is around 12% (Banay-Schwartz et al., 1992). This leaves us with the amount of protein per g of wet weight of the rat brain. Next we divide by the average density of a rat brain which is *ρ_b_* = 1.05*g/ml* (DiResta et al., 1990) to find the amount of receptors per unit of volume of the rat striatum. Finally, we divide by the volume fraction *α*, the fraction of the brain volume that is taken up by the extracellular space in the rat brain, to obtain the receptor concentration of the receptor in the extracellular medium. The procedure ends here for the D1 receptors since there is no evidence that D1 receptors are internalized in the baseline state (Prou et al., 2001). However, a large fraction of the D2 receptors is retained in the endoplasmatic reticulum of the neuron (Prou et al., 2001), reducing the amount of receptors that contribute to the concentration of receptors in the extracellular medium by *f^m^*, the fraction of receptors protruding into the extracellular medium.

In addition to the receptor concentration, the kinetic constants of the receptors are key parameters in our slow kinetics model. In an equilibrium measurement in the canine caudate nucleus the dissociation constant of low affinity DA binding sites, corresponding to D1 receptors (Maeno, 1982), has been measured as *K_D_* = 1.6*μM* (Sano et al., 1979). However, when calculating *K_D_* (using Eq. 8) from the measured kinetic constants (Sano et al., 1979) the value is 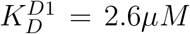. To be more easily comparable to other simulation works (Dreyer et al., 2010) and direct measurements (Richfield et al., 1989; Sano et al., 1979) we choose 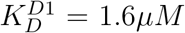 in our simulations. For this purpose we modified both the 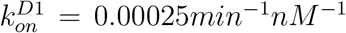 and 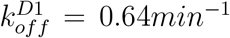 rate measured (Sano et al., 1979) by ~ 25%, making 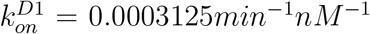 slightly faster and 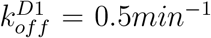 slightly slower, so that the resulting 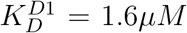. The kinetic constants have been measured at 30°C and are temperature dependent. In biological reactions a temperature change of 10°C is usually associated with a change in reaction rate around a factor of 2-3 (Reyes et al., 2008). However, the conclusions of this paper do not change for an increase in reaction rates by a factor of 2 — 3 (see **Fig. 9**). It should also be noted that the measurements of the commonly referenced *K_D_* (Richfield et al., 1989) have been performed at room temperature.

The kinetic constants for the D2 receptors were obtained from measurements at 37°*C* of high affinity DA binding sites (Burt et al., 1976), which correspond to the D2 receptor (Maeno, 1982). The values are 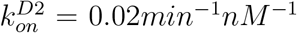 and 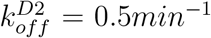, which yields 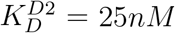, in line with the values measured in (Richfield et al., 1989). As the off-rate of the D1 and D2 receptors 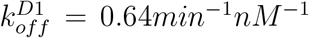 and 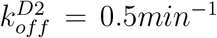 is quite similar, the difference in 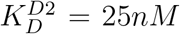 and 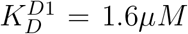 is largely due to differences in the on-rate of the receptors. This is important because the absolute rate of receptor occupancy depends linearly not only on the on-rate, but also on the receptor concentration (see Eq. 5), which means that a slower on-rate could be compensated for by a higher number of receptors.

The parameters used in the simulations are summarized in Tab. 1.

**Table 1:** Receptor parameters

| Measured values |  |  |
| --- | --- | --- |
| Parameter |  | Source |
| $[D1]^m$ in pmol/mg protein | 2840 | (Richfield et al., 1989) |
| $[D2]^m$ in pmol/mg protein | 696 | (Richfield et al., 1989) |
| $\epsilon$ | 0.12 | (Banay-Schwartz et al., 1992) |
| $\alpha$ | 0.2 | (Syková and Nicholson, 2008) |
| $\rho_{brain}$ in g/ml | 1.05 | (DiResta et al., 1990) |
| $f_{D1}^{membrane}$ | 1.0 | (Prou et al., 2001) |
| $f_{D2}^{membrane}$ | 0.2 | (Prou et al., 2001) |
| $k_{on}^{D1,orig}$ in $nm^{-1}min^{-1}$ | 0.00025 | (Sano et al., 1979) |
| $k_{off}^{D1,orig}$ in $min^{-1}$ | 0.64 | (Sano et al., 1979) |
| $k_{on}^{D2}$ in $nm^{-1}min^{-1}$ | 0.02 | (Burt et al., 1976) |
| $k_{off}^{D2}$ in $min^{-1}$ | 0.5 | (Burt et al., 1976) |
| Derived Parameters |  |  |
| Parameter |  | Source |
| $[D1]$ in $nM$ | $\approx 1600$ | Eq.(17) |
| $[D2]$ in $nM$ | $\approx 80$ | Eq.(17) |
| $k_{on}^{D1,used}$ in $nm^{-1}min^{-1}$ | 0.0003125 | see Text |
| $k_{off}^{D1,used}$ in $min^{-1}$ | 0.5 | see Text |

### Dopamine signals

In our model we assumed a baseline [DA] of [*DA*]^tonic^ = 20 *nM* (Dreyer et al., 2010; Dreyer, 2014; Venton et al., 2003; Suaud-Chagny et al., 1992; Borland et al., 2005; Justice Jr, 1993; Atcherley et al., 2015). We modelled changes in [DA] to mimic DA signals observed in experimental studies. We use three types of single pulse DA signals: (long-)bursts, burst-pauses, and ramps.

The (long-)burst signal mimics the effect of a phasic burst in the activity of DA neurons in the SNc, e.g. in response to reward-predicting cues (Pan et al., 2005). The model burst signal consists of a rapid linear [DA] increase (with an amplitude Δ[*DA*] and rise time *t_rise_*) and a subsequent return to baseline. The return to baseline is governed by Michaelis Menten kinetics with appropriate parameters for the dorsal striatum *V_max_* = 4.0 *μMs*^-1^ and *K_m_* = 0.21 *μM* (Bergstrom and Garris, 2003) and the nucleus accumbens *V_max_* = 1.5 *μMs*^-1^ (Dreyer and Hounsgaard, 2013). In our model the removal of DA is assumed to happen without further DA influx into the system (baseline firing resumes when [DA] has returned to its baseline value). Unless stated otherwise, the long-burst signals are used with a Δ[*DA*] = 200 *nM* and a rise time of *t_rise_* = 0.2 s at *V_max_* = 1.5 *μMs*^-1^, similar to biologically realistic transient signals (Cheer et al., 2007; Robinson et al., 2001; Day et al., 2007).

The burst-pause signal has two components, an initial short, small amplitude burst (Δ[*DA*] = 100 *nM*, *t_rise_* = 0.1 *s*), with the corresponding [DA] returning then to baseline (as for the long burst above). However, there is a second component in the DA signal, in which [DA] falls below baseline, simulating the effect of a pause in DA neuron firing. The length of this firing pause is characterized by the parameter *t_pause_*. This burst-pause [DA] signal reflects the DA cell firing pattern consisting of a brief burst followed by a pause in activity (Pan et al., 2008; Schultz, 2016).

The ramp DA signal is characterized by the same parameters as the burst pattern, but with a longer *t_rise_*, and a smaller Δ[*DA*] (parameter settings provided in each simulation).

For the simulations comparing the area under the curve of the input DA signal with the resulting receptor occupancy (**Fig. 5**) we used the burst, burst-pause, and ramp signals described above with a range of parameter settings. For the burst DA signal we used amplitudes Δ[*DA*]^*max*^ of 50, 100, 150, 200, 250, 300, 350, 400, 500, 600, 700, 800, 900, and 1000 nM. For the ramping DA signals we used rise times *t_rise_* of 0.2, 0.5, 1.0, 1.5, 2.0, 3.0, 4.0, 5.0, 6.0, and 7.0 s, For the burst-pause DA signal we different values for *V_max_* of 1.0, 1.5, 2.0, 2.5, 3.0, 2.5, and 4.0 *μMs*^-1^.

### Behavioural task simulation

To determine whether DA receptor occupancy can integrate reward signals over minutes, we simulated experiments consisting of a sequence of 50 trials. In each sequence the reward probability was fixed. The trials contained either a (long-)burst DA signal (mimicking a reward) or a burst-pause DA signal (mimicking no reward) at the beginning of the trial according to the reward probability of the sequence. The inter-trial interval was 15 ± 5s (**Fig. 8**). We choose this highly simplistic scenario to mimic DA signals in a behavioural task in which the animal is rewarded for correct performance. However, here the specifics of the task are not relevant as our model addresses the integration of the DA receptor occupancy over time. Although we chose to use the burst-pause type signal as shown in **Fig. 2a** as a non-rewarding event, the difference to a non-signal are minimal after the end of the pause (**Figs. 3 and 4**). Each sequence started from a baseline receptor occupancy, assuming a break between sequences long enough for the receptors to return to baseline occupancy (around 5 minutes). For the simulations shown in **Fig. 4** all trials started exactly 15 s apart.

While for the simulations shown in **Fig. 4** the sequence of DA signals was fixed, we also simulated a behavioural task with stochastic rewards (**Fig. 8**). There we simulated reward probabilities from 0% to 100% in 10% steps. For each reward probability we ran 500 sequences, and calculated the mean receptor occupancy over time (single realizations shown in **Fig. 8a, c**). To investigate whether the receptor occupancy distinguished between different reward probabilities we applied a simple classifier to the receptor occupancy time course.

The classifier was used to compare two different reward probabilities at a time. At each time point during the simulated experiment it was applied to a pair of receptor occupancies, e.g. one belonging to a 50% and one to a 30% reward probability sequence. For each sequence the classifier assigned the current receptor occupancy to the higher or lower reward probability depending on which reward probabilities mean (over 500 sequences) receptor occupancy was closer to the current receptor occupancy. As we knew the underlying reward probability of each sequence we were able to calculate the true and false positive rates and accuracy for each time point in our set of 500 sequences for both the D1R and D2R (**Fig. 8e, f**). The accuracy was calculated based on all time points between 200 and 800s within a sequence to avoid the effect of the initial “swing-in” and post-sequence DA levels returning to baseline.

## Results

Before investigating the role of the receptor kinetics in response to different DA signals, we started by establishing the receptor binding at baseline [DA], taking into account the different numbers of D1 and D2 receptors in the striatum. For a stable baseline [DA] the receptor affinities can be used to calculate receptor occupation (see Methods, Eq. 1).

First, we investigated receptor binding for a range of affinities (**Fig. 1**), reflecting the range of measured values in different experimental studies (Neve and Neve, 1997). We report the resulting receptor occupancy in terms of the concentration of D1Rs and D2Rs bound to DA (denoted as [*D*1 — *DA*] and [*D*2 — *DA*], respectively). Due to the low affinity of D1Rs, at low baseline [DA] only a small fraction of D1 receptors may be occupied. However, there are overall more D1Rs than D2Rs (Richfield et al., 1989), and ≈ 80% of D2Rs are retained in the endoplasmatic reticulum (Prou et al., 2001). Therefore, the concentration of D1Rs in the membrane available to extracellular DA is a lot higher than the concentration of D2Rs (e.g. 20 times more in the nucleus accumbens; Nishikori et al., 1980; see Methods). Thus, in our simulation, the actual concentration of bound D1Rs ([*D*1 — *DA*] ≈ 20nM) was, at DA baseline, much closer to the concentration of bound D2Rs ([*D*2 — *DA*] ≈ 35nM) than suggested by the different D1 and D2 affinities alone. We further confirmed that this was not due to a specific choice of the dissociation constants in the model, as [*D*1 — *DA*] and [*D*2 — *DA*] remained similar over the range of experimentally measured D1R and D2R affinities (Neve and Neve, 1997) (**Fig. 1a**). This suggests that [*D*1 — *DA*] is at most twice as high as [*D*2 — *DA*] instead of 40 times higher as suggested by the difference in fraction of bound receptors. Therefore, [*D*1 — *DA*] and [*D*2 — *DA*] might be better indicators for the signal transmitted to MSNs, as the fraction of bound receptors neglects the different receptor type abundances.

**Figure 1:**
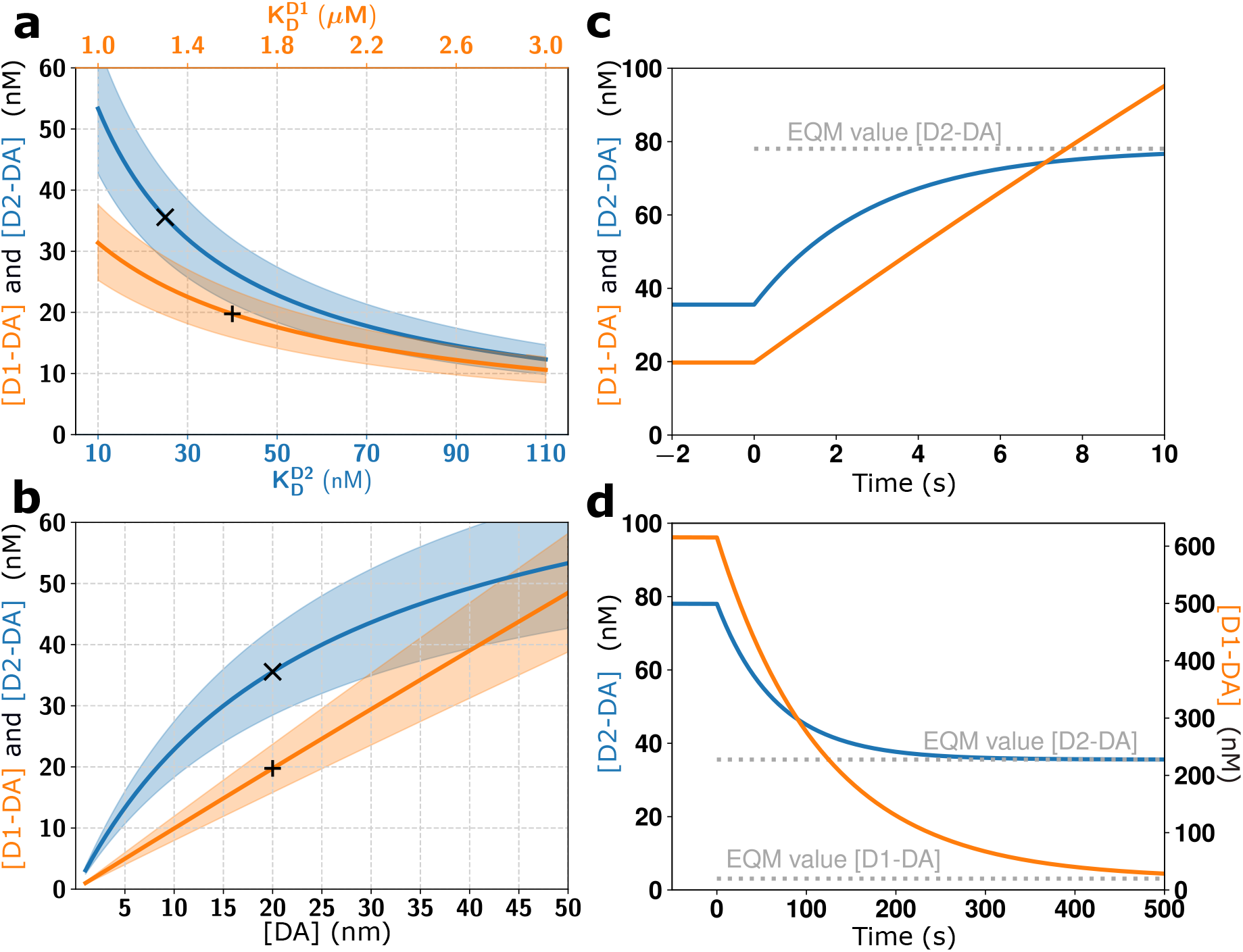
Baseline levels of D1 and D2 receptor occupation and impact of slow kinetics. **(a)** Equilibrium values of absolute concentration of receptors bound to DA as a function of receptor affinities. Here, baseline [DA] was fixed at 20 nM. **(b)** Equilibrium values of absolute concentration of receptors bound to DA as a function of baseline [DA]. Here 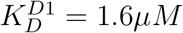 and 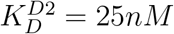. ‘×’ and +’ indicate the model default parameters. Coloured bands mark the range of values for up to ±20% different receptor abundances. **(c)** Temporal dynamics of D1 and D2 receptor occupancy for a large step up from [*DA*] = 20*nM* to [*DA*] = 1*μM*. The gray dotted line shows that equilibrium value (EQM). **(d)** Same as in **c** but for a step down from [*DA*] = 1*μM* to [*DA*] = 20*nM*.

Next, we investigated the effect of slow [DA] changes (Grace, 1995; Schultz, 1998; Floresco et al., 2003) by exposing our model to changes in the [DA] baseline. For signalling timescales that are long with respect to the half-life time of the receptors (*t_s1ow_* >> *t*_1/2_ ≈ 80*s*), we used the dissociation constant to calculate the steady state receptor occupancy. We found that for a range of [DA] baselines (mimicking slow changes in [DA]), there was less than two-fold difference between [*D*1 — *DA*] and [*D*2 — *DA*] (**Fig. 1b**), because of the different abundances of D1 and D2 receptors. This is in contrast to affinity-based models, which suggest that D2Rs are better suited to encode slow or tonic changes in [DA]. Interestingly the change of [*D*1 — *DA*] was almost linear in [DA], while the change of [*D*2 — *DA*] showed nonlinear effects due to the change in available free D2R. Thus, based on these results, it could even be argued that D1Rs are better at detecting tonic signals at high [DA] levels, since they do not saturate as easily.

While for baseline and slow changes in [DA] the receptor occupation can be determined based on the receptor affinity, fast changes in [DA] also require a description of the underlying receptor kinetics. To investigate the effect of typical DA signals on receptor occupation, we developed a kinetics model incorporating binding and unbinding rates that determine the overall receptor affinity (see Methods, Eq.8, 9). The available experimental measurements indicate that the different D1R and D2R affinities are largely due to different binding rates, while their unbinding rates are similar (Burt et al., 1976; Sano et al., 1979; Maeno, 1982; Richfield et al., 1989). We incorporated these measurements into our kinetics model and investigated the model’s response to a variety of fast DA signals.

We started by measuring the model response to a [DA] step change from 20nM to 1*μM*. This is quite a large change compared to phasic DA signals *in vivo* (Robinson et al., 2001; Cheer et al., 2007; Hamid et al., 2016), which we choose to illustrate that our results are not just due to a small amplitude DA signal. We found that binding to both receptor subtypes increased very slowly. Even for the high affinity D2Rs it took more than 5s to reach their new equilibrium (**Fig. 1c**). Thus, unlike the affinity-based model, our model suggests that the D2Rs will not saturate for single reward events, which last overall for up to ≈ 3s. Note that the non-saturation is independent of the abundance of the receptors and is only determined by the kinetics of the receptors (see Methods). Due to their slow unbinding, D1Rs and D2Rs also took a long time to return to baseline receptor occupancy after a step down from [*DA*] = 1*μM* to [*DA*] = 20*nM* (**Fig. 1d**). Thus, we conclude that with slow kinetics of receptor binding both D1Rs and D2Rs can detect single phasic DA signals and that both remain occupied long after a high [DA] has returned to baseline.

### DA receptor binding kinetics for different types of DA signals

Next, we investigated [*D*1 — *DA*] and [*D*2 — *DA*] for three different types of DA signals (**Fig. 2**). The first signal was a phasic DA increase (‘long burst’, **Fig. 2a**), mimicking responses to rewards and reward-predicting stimuli (Robinson et al., 2001; Cheer et al., 2007). The second signal was a brief phasic DA increase, followed by a decrease (‘burst-pause’, **Fig. 2a**), mimicking responses to conditioned stimuli during extinction (Pan et al., 2008) or to other salient stimuli (Schultz, 2016). The third signal was a prolonged DA ramp, mimicking a value signal when approaching a goal (Howe et al., 2013; Hamid et al., 2016) (**Fig. 2b**). In the affinity-based model with instant kinetics the D1Rs mirrored the [DA] time course for all three types of signals, since even at [*DA*] = 200*nM* D1Rs are far from saturation. By contrast, D2Rs showed saturation effects as soon as 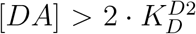, leading to differing D1 and D2 time courses (**Fig. 2**, middle and bottom rows, grey traces). Importantly, in our model with slow kinetics, the time courses of [*D*1 — *DA*] and [*D*2 — *DA*] were nearly identical, supporting that both receptor types are equally affected by phasic DA signals. This was the case for all the three signals: burst, burst-pause and ramping DA signals. The only difference between the [*D*1 — *DA*] and [*D*2 — *DA*] time courses were the absolute amplitudes. For example, [*D*2 — *DA*] started from a baseline about twice as high as [*D*1 — *DA*], but then also responded to the long burst DA signal with a change about twice as high. The similarity of [*D*1 — *DA*] and [*D*2 — *DA*] responses to both slow (**Fig. 1b**) and fast (**Fig. 2**) [DA] changes indicates that the different DA receptor types respond similarly independent of the timescale of [DA] changes. It could even be argued that D2Rs are better at detecting phasic DA signals, since they respond with a larger absolute change in occupied receptors.

**Figure 2:**
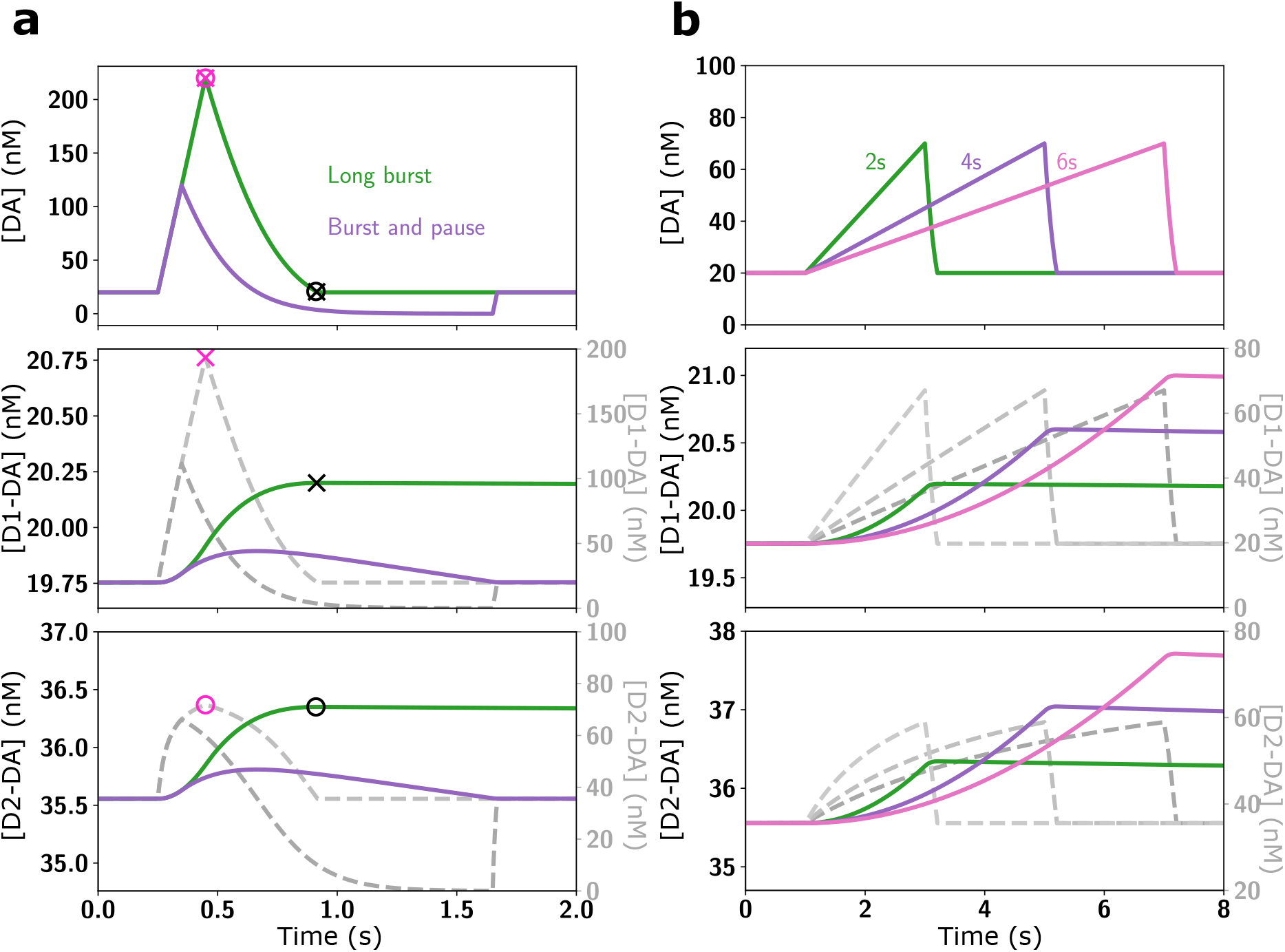
Impact of receptor kinetics on responses to different DA signals. **(a)** Two different DA signals, long burst and burst-pause, were simulated. The top panel shows the time course of the model [DA] input signal, with the resulting changes in D1 and D2 receptor occupancy in the middle and bottom panel, respectively. The colored traces in the middle and bottom panel show the resulting [DA-D1] and [DA-D2] for the realistic kinetics model (left scales), and the dashed gray traces show the corresponding values for the affinity-based model (right scales). The timing of the maximum receptor occupancy (‘×’ and ‘o’ for D1 and D2, respectively) coincides for instant kinetics (purple symbols) with the [DA] peak (combined x and o in top panel), while for slow kinetics (black symbols) it coincides with the offset of the [DA] signal instead (combined x and o in top panel). **(b)** Same as in (a) but for ramping DA signals.

To understand why the D1Rs and D2Rs respond in a similar fashion, we considered the relevant model parameters in more detail. The binding rate constants of D1Rs and D2Rs differ by a factor of ≈ 60 (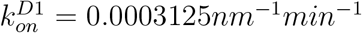 and 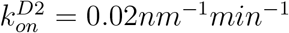; Burt et al., 1976; Sano et al., 1979; Maeno, 1982; see also Methods), suggesting faster D2Rs. However, experimental data suggests that there are ≈ 40 fold more unoccupied D1 receptors ([*D*1] ≈ 1600*nM*) than unoccupied D2 receptors ([*D*2] ≈ 40*nM*) on MSN membranes in the extracellular space of the rat striatum (Nishikori et al., 1980). Therefore, the absolute binding rate 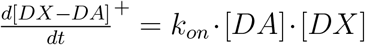 differs only by a factor of ≈ 1.5 between the D1Rs and D2Rs. That is, the difference in the kinetics of D1Rs and D2Rs is compensated by the different receptor numbers, resulting in nearly indistinguishable aggregate kinetics (**Fig. 2**). This is consistent with recent experimental findings which showed that D2R expressing MSNs can detect phasic [DA] signals (Marcott et al., 2014; Yapo et al., 2017).

The dynamics introduced by the slow kinetics in our model also affected the time course of DA signalling. With instant kinetics the maximum receptor occupancy was reached at the peak [DA] (**Fig. 2**, middle and bottom rows). By contrast, for slow kinetics the maximum receptor occupancy was reached when [DA] returned to its baseline (**Fig. 2a**) because as long as [DA] was higher than the equilibrium value of [D1-DA] and [D2-DA], more receptors continued to become occupied. Therefore, for all DA signals, the maximum receptor occupancy was reached towards the end of the pulse.

Another striking effect of incorporating receptor kinetics was that a phasic increase in [DA] kept the receptors occupied for a long time (**Fig. 2a** green traces). However, when a phasic increase was followed by a decrease, [*D*1—*DA*] and [*D*2—*DA*] returned to baseline much faster (**Fig. 2a** purple traces). This indicates that burst-pause firing patterns can be distinguished from pure burst firing patterns on the level of the MSN DA receptor occupancy. This supports the view that the fast component of the DA firing patterns (Schultz, 2016) is a salience response, and points to the intriguing possibility that the pause following the burst can, at least partly, revoke the receptor-ligand binding induced by the burst. In fact, for each given burst amplitude, a sufficiently long pause duration can cancel the receptor activation (**Fig. 3**), with larger [DA] amplitudes requiring longer pauses to cancel the activation. Thereby, the burst-pause firing pattern of DA neurons could effectively signal a reward “false-alarm”.

**Figure 3:**
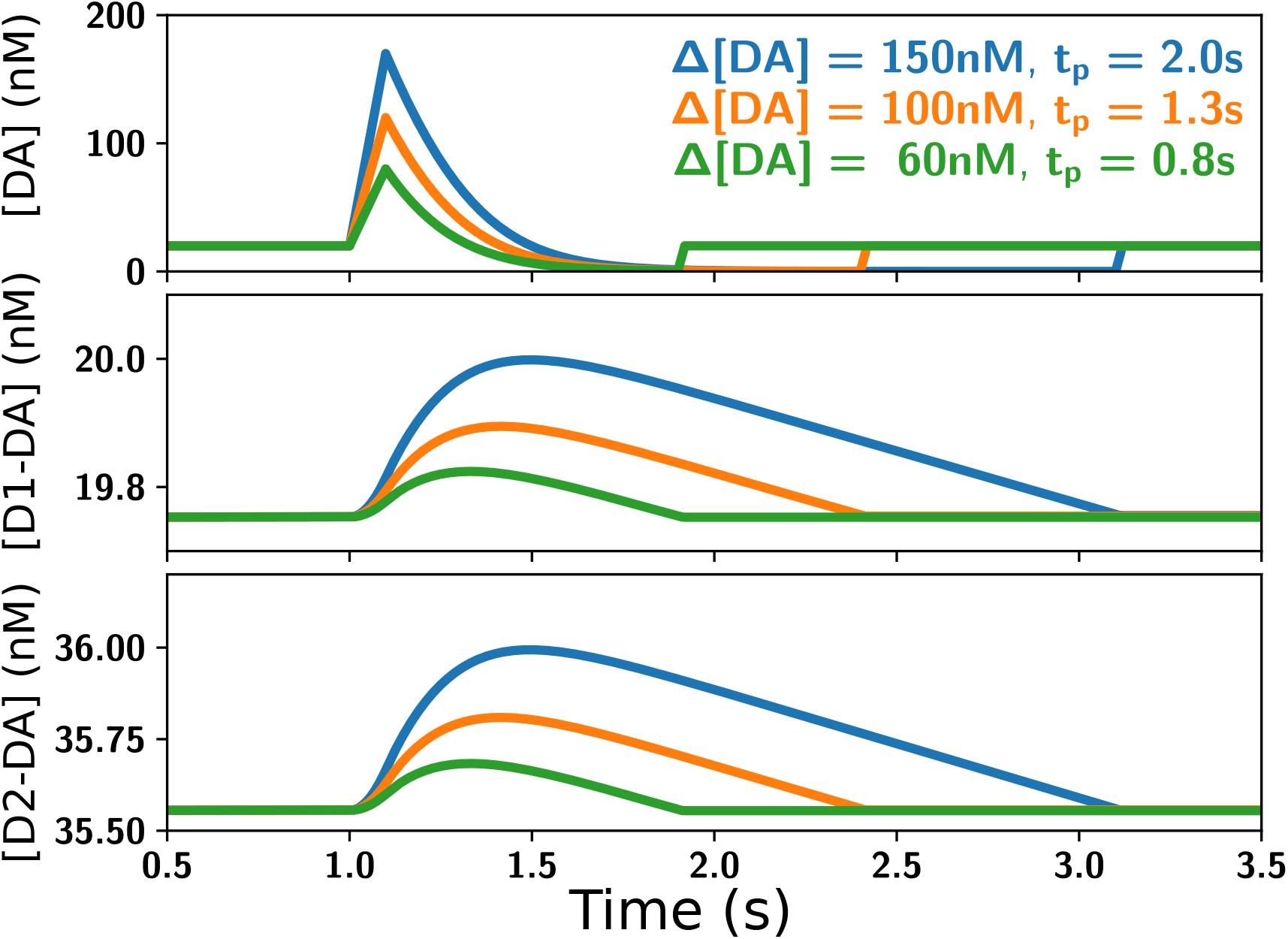
Burst-pause DA signals (top panel) did not lead to a prolonged D1 or D2 receptor occupation (middle and bottom panels, respectively). The initial increase in receptor occupation due to the burst component was quickly cancelled by the unbinding that occurred during the pause component. Higher burst amplitudes required a longer pause duration for the cancellation of the receptor occupation.

The long time it took [*D*1 — *DA*] and [*D*2 — *DA*] to return to baseline after phase DA signals (**Fig. 2a**) indicates that the receptor occupation integrates DA signals over time. To examine this property, we simulated a sequence of DA signals on a timescale relevant for behavioural experiments (**Fig. 4**). Each sequence consisted of 50 events and each event was separated by 15 s. Three different types of sequences were tested: 50 phasic DA bursts, 40 phasic DA bursts followed by 10 burst-pause signals, and 40 phasic DA bursts followed by 10 non-events. We found that both [*D*1 — *DA*] and [*D*2 — *DA*] accumulated over the sequence of DA signals. The sawtooth shape of the curves was due to the initial unbinding of the receptors, which was then interrupted by the next DA signal 15 s later. At higher levels of receptor activation, the amount of additional activated receptor per DA pulse was reduced since there are less free receptors available, and the amount of receptors unbinding during the pulse duration was increased because more receptors were occupied. The accumulation occurred as long as the time interval between the DA signals was shorter than ≈ 2 · *t*_1/2_. Together, the shape and period of the DA pulses lead to the formation of an equilibrium, visible here as a plateau for the absolute amount of occupied receptor. This occurred at the level at which the amount of receptors unbinding until the next DA burst was the same as the amount of receptors getting occupied by the DA burst. Finally, the burst-pause events did not lead to an accumulation of occupied receptors over time. In fact, the receptor occupation was the same for burst-pause and non-event, except during the short burst component of the burst-pause events (note the overlapping green and orange curves in **Fig. 4**). This extends the property of burst-pause signals as “false alarm” signals to a wide range of occupancy levels.

**Figure 4:**
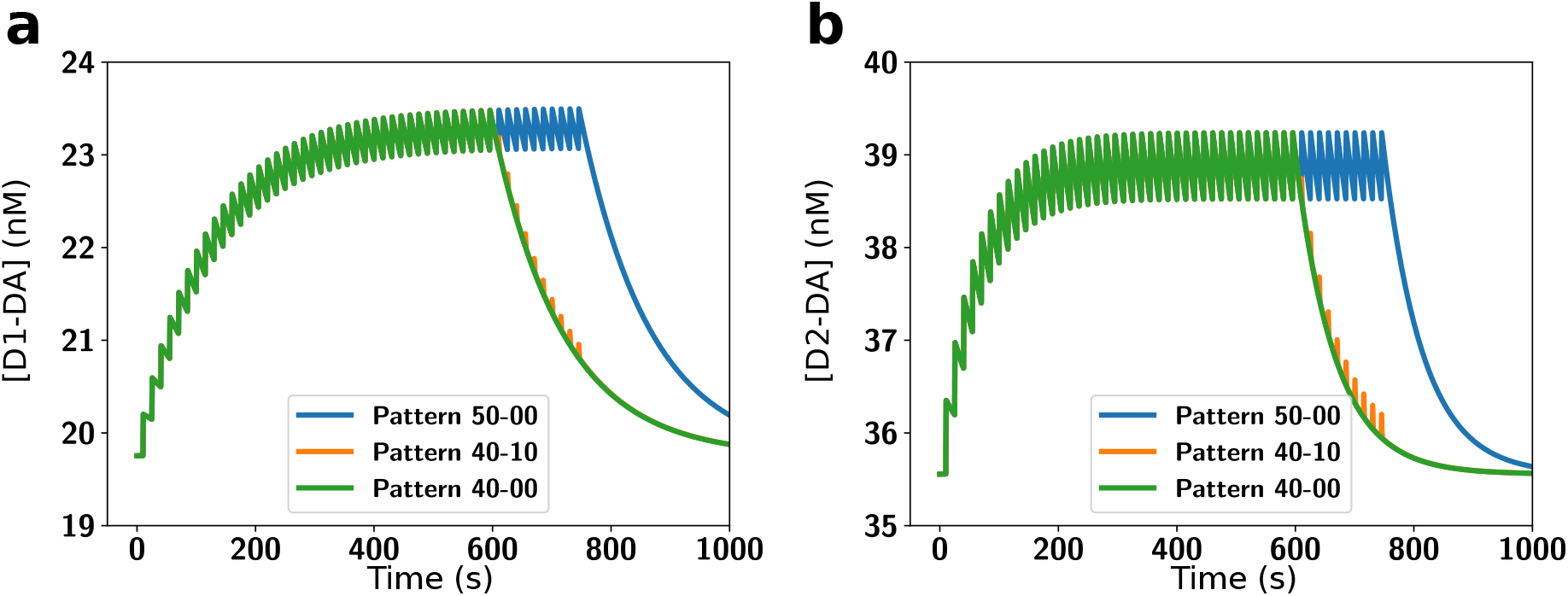
D1 and D2 receptor occupation integrates DA signals over a behavioural time scale. **(a)** The absolute receptor occupancy for D1Rs for three different types of sequences consisting of 50 DA events each. The sequences consisted of 50 long burst events (blue), 40 long burst followed by 10 burst-pause events (orange) and 40 long burst events followed by 10 non-events (green, for comparison). **(b)** Same as in a but for D2Rs. Note that for the time course of the overall receptor occupation burst-pause signals are basically identical to non-events.

Incorporating the slow kinetics in the model is crucial for functional considerations of the DA system. Currently, following the affinity-based model, the amplitude of a DA signal (i.e. peak [DA]) is often considered as a key signal, e.g. in the context of reward magnitude or probability (Morris et al., 2004; Tobler et al., 2005; Hamid et al., 2016). However, as DA unbinds slowly (over tens of seconds; **Fig. 1d**) and the binding rate changes approximately linearly with [DA], the amount of receptor occupancy does not primarily depend on the amplitude of the [DA] signal. Due to the linearity of the binding rate, the receptor occupation increases linearly with time and [DA] – [DA]_baseiline_, while the unbinding is negligible as long as *t* << *t*_1/2_. Therefore the integral of the [DA] time course should be a close approximation of the receptor occupation for signals that are shorter than the half-life time of the receptors. We confirmed this consideration by simulating a range of DA signals (burst, burst-pauses, and ramps) with different durations and amplitudes. For each DA signal we compared its area under the curve with the resulting peak change in the absolute receptor occupancy. For both D1R and D2R we found that the maximum receptor activation was proportional to the area under the curve of the [DA] signal, while independent of its specific time course (**Fig. 5**). The small deviation from the proportionality seen for large-area DA signals for the D2Rs (**Fig. 5b**) was due to the decrease in the amount of free receptor as more and more receptors were bound. In this regime the assumption that the binding rate is linear with [DA] was slightly violated leading to the non-proportionality. The overall striking proportionality of the integral of the DA signal with receptor binding indicates that D1Rs and D2Rs act as slow integrators of the DA signal. Interestingly, this means that DA ramps, even with a relatively small amplitude (**Fig. 2b** and **Fig. 7**), are an effective signal to occupy DA receptors. In contrast, for locally very high [DA] (e.g. at corticostriatal synapses during phasic DA cell activity; Grace et al., 2007) our model predicts that the high concentration gradient would only lead to a very short duration of this local DA peak and thereby make it less effective in occupying DA receptors.

**Figure 5:**
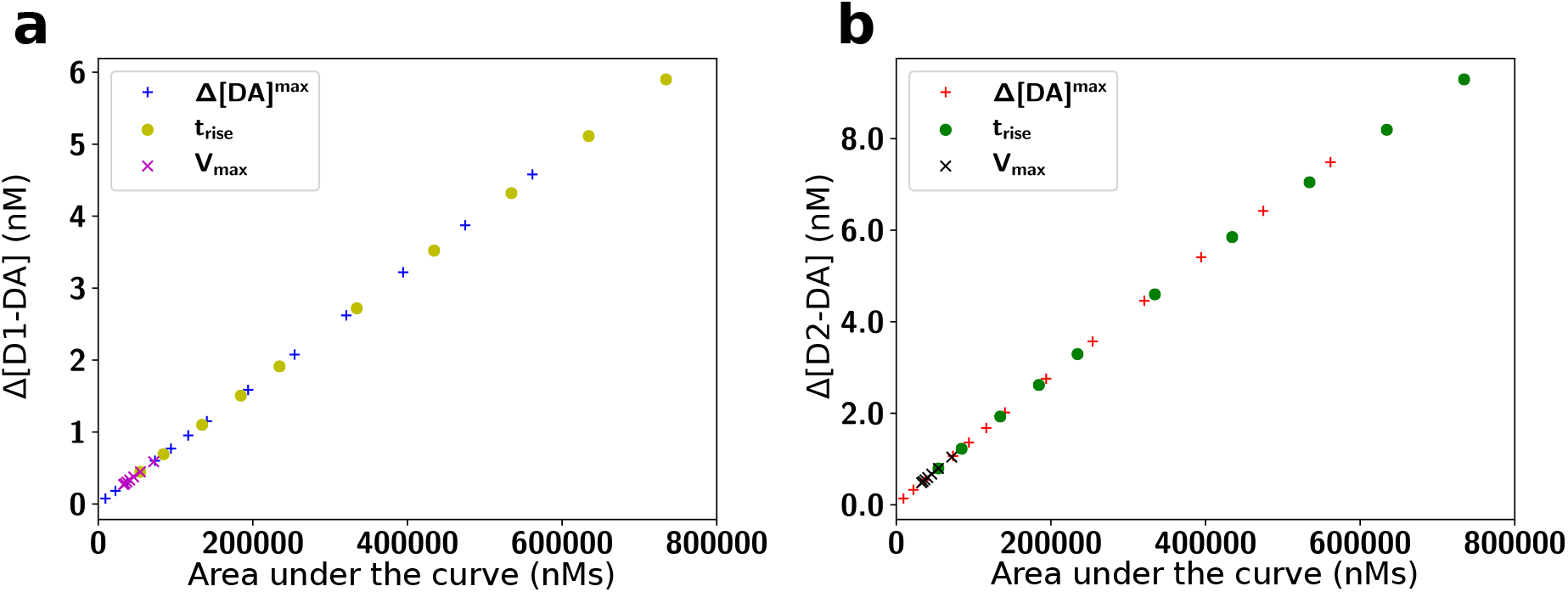
DA receptor occupation is proportional to the area under the curve of DA signals. The peak change in absolute receptor occupancy of D1Rs **(a)** and D2Rs **(b)** shown on the y-axis increased linearly with the area under the curve of the DA pulses. Each data point provides the result of a single simulation with a given parameter setting for the burst amplitude (Δ[*DA*]^*max*^), ramp rise time (*t_rise_*), and DA re-uptake rate (*V_max_*).

To further test the generality of our findings, we examined our model responses systematically for a set of different DA time courses (**Fig. 6** and **Fig. 7**). While the shape of the DA pulses strongly affected the time courses of the receptor activation, D1 and D2 receptor activation were virtually identical for a given pulse shape. For DA bursts with different amplitudes (**Fig. 6a**), higher amplitudes of the DA burst lead to stronger receptor activation. However, the relationship between burst amplitude and receptor occupation was not linear, but instead reflected the area under the curve of the DA pulse (see above). Importantly, despite ‘slow’ kinetics, the onset of the increase in [*D*1 — *DA*] and [*D*2 — *DA*] was immediate and reached relevant levels in a nanomolar range within a few 100 ms. For a fixed burst amplitude, we also determined the effect of different DA re-uptake rates to look at potential differences in DA signalling in dorsal and ventral striatum, with fast and slow re-uptake, respectively. This was done by changing the parameter *V_max_* (see Methods), which controlled the time the [DA] took to return to the baseline from the peak value (**Fig. 6b**). While this had only a small visible effect on the input DA signal (**Fig. 6b**, top panel), the resulting [*D*1 — *DA*] and [*D*2 — *DA*] were quite different. This is important because this property is not seen in the affinity-based model, in which the time course of [*D*1 — *DA*] and [*D*2 — *DA*] would simply follow the input [DA] signal.

**Figure 6:**
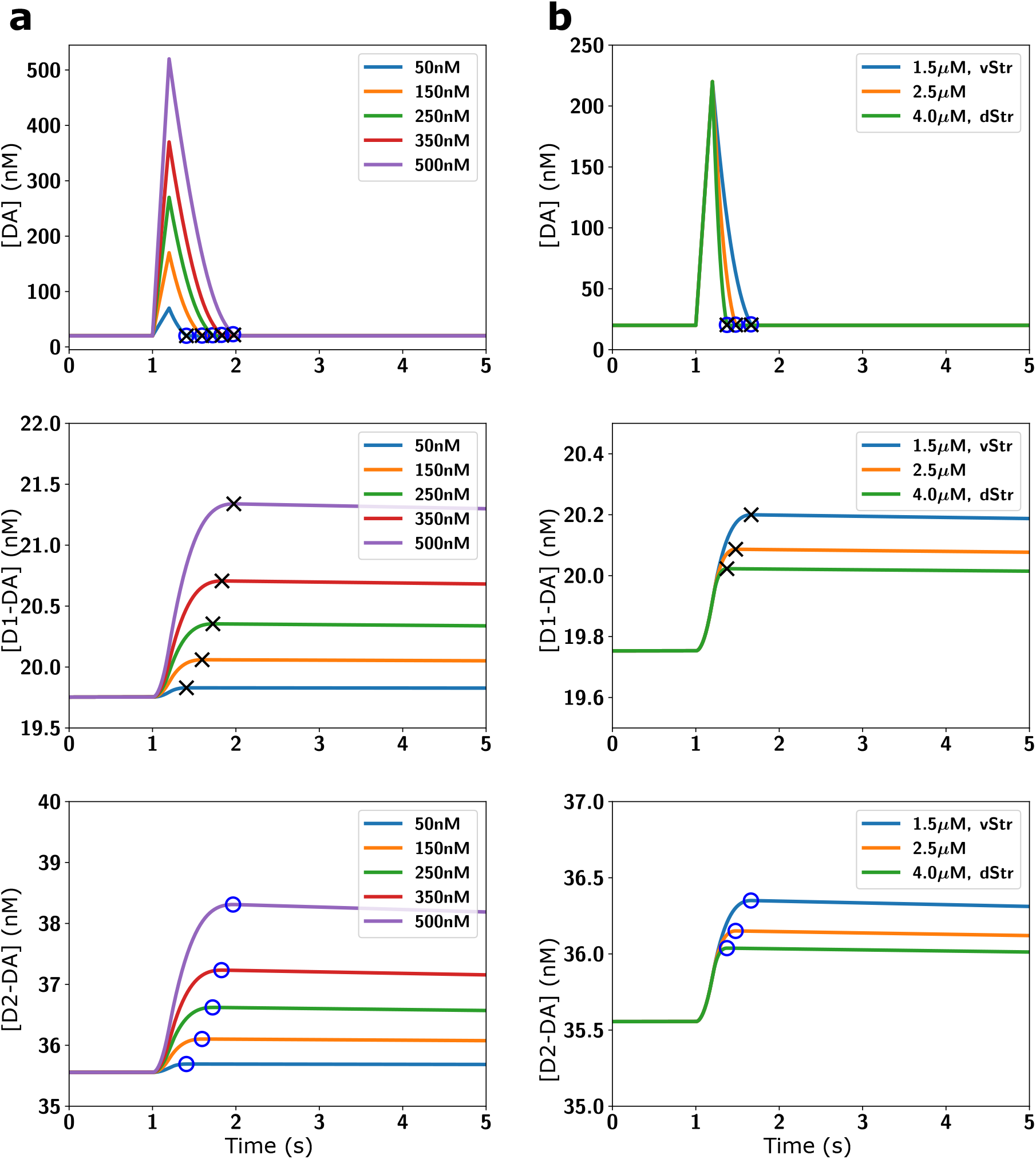
Parameter exploration for phasic DA bursts (top row) with the resulting changes in D1 (middle row) and D2 (bottom row) receptor occupancy. **(a)** Effect of variations in the amplitude Δ[*DA*]^*max*^ of the phasic DA burst (top row) on the D1 (middle row) and D2 (bottom row) receptor occupancy, **(b)** Effect of change in the re-uptake rate *V_max_* rate (top row) on the D1 (middle row) and D2 (bottom row) receptor occupancy. *V_max_* was changed to mimic conditions for the ventral and dorsal striatum. Blue circles and black crosses mark the time points of maximum receptor occupancy for D1 and D2, respectively. Note that for both D1R and D2R the time of maximum receptor occupancy was near the end of the DA signal and that D1Rs and D2Rs behaved similarly independent of the specific parameters of the DA pulse.

Next, we examined DA ramps with different time courses, but the same maximal amplitude. Again, consistent with our consideration of the important role of the area under the curve of DA signals, we found that longer ramps lead to larger DA receptor occupation (**Fig. 7a**). We investigated the DA signals that included the effects of pauses in DA cell activity further. First, we tested burst-pause signals and determined the role of the duration of the pause. For a fixed burst amplitude and duration, a different duration of the subsequent pause lead to differing receptor activation levels when the burst-pause signal was over (**Fig. 7b**). This indicates that DA pauses are very effective in driving the receptor occupation quickly back to baseline (i.e. within few seconds) because, in this case, the receptor occupation changes reflect solely the unbinding rates. In contrast, for a burst followed by a return to baseline [DA], the decrease in receptor occupation would be slower because during the baseline portion of the signal both binding and unbinding processes play a role. In this case the binding counteracts some of the unbinding. In this context we also looked a pure DA pauses (i.e. without a preceding burst), e.g. reflecting DA cell responses to aversive stimuli (Schultz, 2007) that lead to reductions in [DA] (Roitman et al., 2008). These signals also lead to fast decreases in [D1-DA] and [D2-DA], with the duration of the pause having a strong effect on the amplitude and duration of the decrease (**Fig. 7c**).

**Figure 7:**
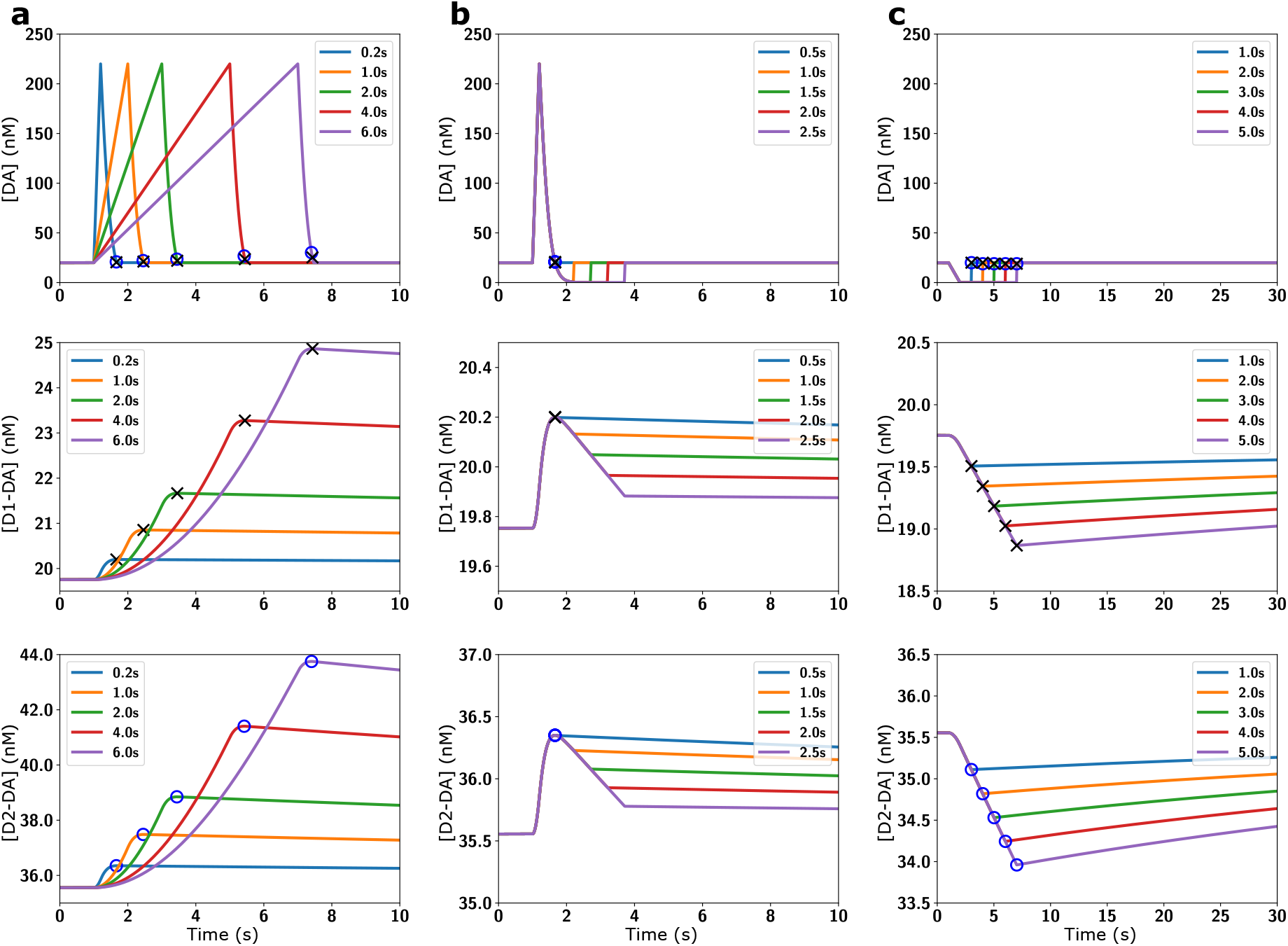
Parameter exploration for different DA signals (top row) with the resulting changes in D1 (middle row) and D2 (bottom row) receptor occupancy. **(a)** D1 (middle row) and D2 (bottom row) receptor occupancy for different rise time *t_rise_* of the DA ramps (top row). The rise time controls the amount and duration of D1 (middle row) and D2 (bottom row) receptor occupancy. **(b)** D1 (middle row) and D2 (bottom row) receptor occupancy for different pause duration *t_pause_* of the burst-pause type DA signals (top row). **(c)** D1 (middle row) and D2 (bottom row) receptor occupancy for different pause duration *t_pause_* of DA pauses (without a preceding burst). Such a DA pause led to a fast reduction of receptor occupancy, which took 10s of seconds to return to baseline. The blue circles and black crosses mark the time points of maximum receptor occupancy for D1 and D2, respectively (a-b), or of minimal receptor activation (c). Note that for both D1R and D2R the time of maximum (or minimum for c) receptor occupancy was near the end of the DA signal and that D1Rs and D2Rs behaved similarly independent of the specific parameters of the DA pulse.

### D1R and D2R occupancy in a probabilistic reward task paradigm

A general effect of the slow kinetics was that DA receptors remained occupied long after the DA pulse was over (**Fig. 2**), so that the effect of DA pulses was integrated over time (**Fig. 4**). To investigate the information that is preserved in the receptor occupation about DA signals on time scales relevant for behavioural tasks, we simulated sequences with probabilistic DA events (see Methods). First, we compared sequences, in which every 15 ± 5s there was a DA burst with either 30%, 50%, or 70% probability (**Fig. 8a, c**). The resulting changes in [*D*1 — *DA*] and [*D*2 — *DA*] confirmed the integration of DA pulses over minutes. The integration of DA bursts was due to DA bursts arriving before the receptor occupation caused by the previous pulses had decayed, leading to an increased receptor activation compared to single DA bursts (**Fig. 4**). We then examined whether the DA receptor occupancy can distinguish different reward probabilities by using a simple classifier comparing two sequences with each other (see Methods). We tested sequences from 0% to 100% probability in steps of 10%, and ordered the resulting classification success in terms of the difference in reward probability between the two sequences (**Fig. 8b, d, e, f**). For example, a comparison between a 30% and a 70% reward probability sequence yields a data point for a 40% difference. For both D1 and D2 receptors, we found that already for differences of 10% the classification exceeded chance level, and yielded near perfect classification around a 40% difference. Overall, the classification was slightly better for D1R due to their slower unbinding rate and more stable plateau response (**Fig. 4**). The successful classification of reward probabilities demonstrates that it would be possible for striatal neurons to read out different reward rates from DA receptor occupancy in a behavioural task. This provides a potential neural substrate for how fast DA signals can lead to an encoding of the slower reward rate, which can be utilized as a motivational signal (Mohebi et al., 2019).

**Figure 8:**
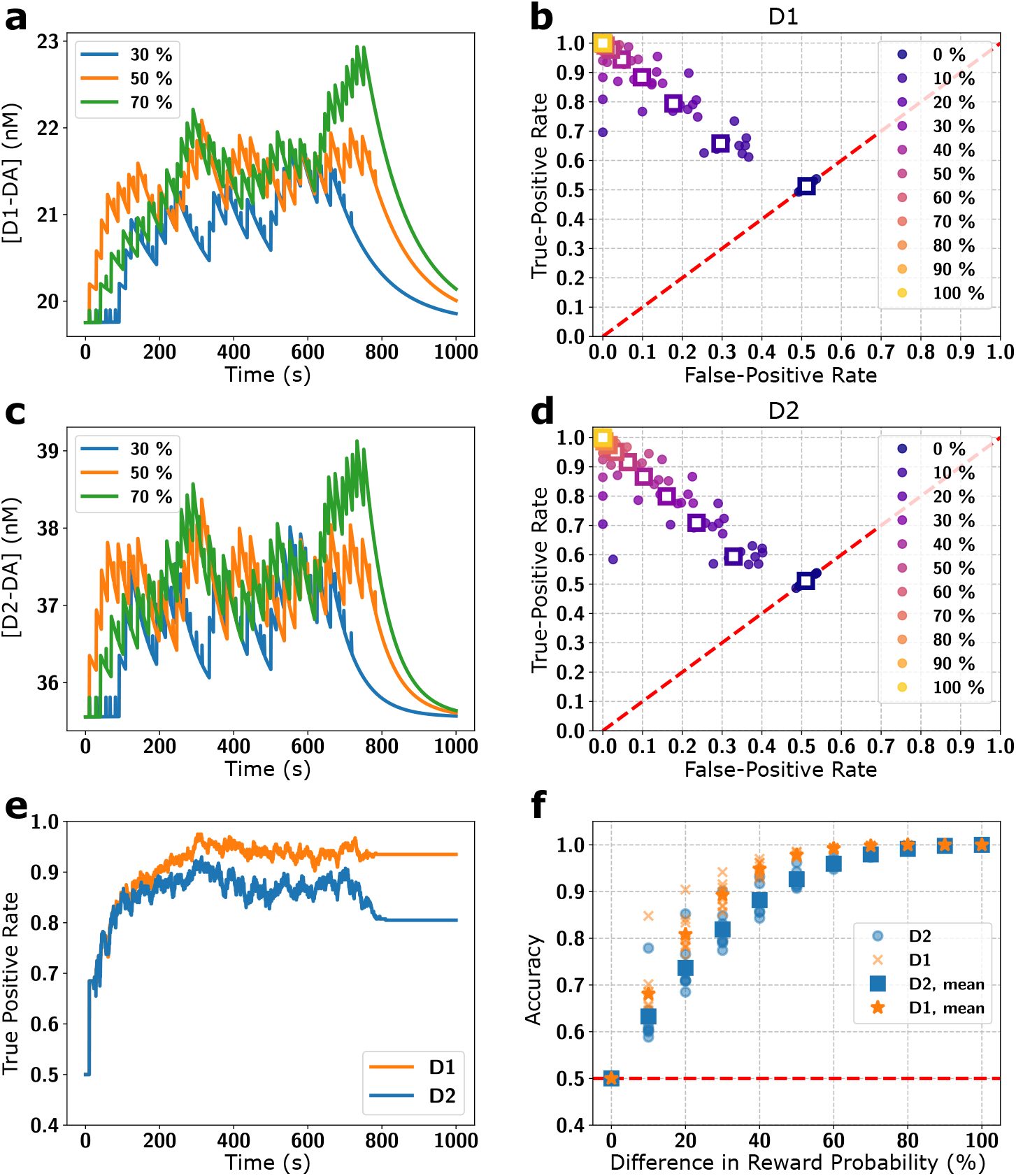
Encoding of reward rate by integration of DA signals over minutes in a simulation of a behavioural task. **(a)** Time course of D1 receptor occupancy for sequences of 50 trials with a reward probability, as indicated, in each trial. **(b)** True and false positive rates of the difference in reward probability based on the D1 and D2 receptor occupancy by a simple classifier. Each dot indicates the true and false positive rate from a simulation scenario with the difference in reward probability indicated by the colour. The colour indicates the difference in reward probability (e.g. a 10% difference in purple occurs for 80% vs. 90%, 70% vs. 80%, etc.), and the squares denote the corresponding averages. The red line indicates chance level performance, and a perfect classifier would be at 1.0 true and 0.0 false positive rate. **(c, d)** The same as in panels a and b but for D2 receptors. **(e)** True positive rates for the classification in a sample session (70% vs 30% reward probability) based on the receptor occupancy of D1 (orange) and D2 (blue) receptors. After a short “swing-in” the receptors distinguished between a 70% and a 30% reward rate. **(f)** Accuracy of the classifier for a range of reward probability differences for the D1 (orange) and D2 (blue) receptors for individual sessions and corresponding session averages.

### Validation for fast binding kinetics

Our model assumption of slow kinetics was based on neurochemical estimates of wildtype DA receptors (Burt et al., 1976; Sano et al., 1979; Maeno, 1982). In contrast, recently developed genetically-modified DA receptors, used to probe [DA] changes, have fast kinetics (Sun et al., 2018; Patriarchi et al., 2018). Although the kinetics of the genetically modified DA receptors are unlikely to reflect the kinetics of the wildtype receptors (see Discussion), we also examined the effect of faster DA kinetics in our model. Fast kinetics were implemented by multiplying *k_on_* and *k_off_* by a factor *q*, keeping *K_D_* constant. We found that the similarity between [*D*1 — *DA*] and [*D*2 — *DA*] persists even if the actual kinetics were a 100 times faster than assumed in our model (**Fig. 9**). This was the case for all types of [DA] signals because the difference between the aggregate D1 and D2 binding rates (Eq. 5) still only differed by a factor of 1.5. Furthermore, the D2Rs did not show visible saturation effects even for *q* = 100. Faster kinetics mostly affected the amplitude of the receptor response and the time it took to return to baseline receptor occupancy. However, only for *q* = 100 the pauses dropped slightly below baseline receptor occupancy (**Fig. 9a, b**). On a longer time scale with repetitive DA bursts (**Fig. 9e, f**) D1Rs and D2Rs integrated the DA bursts over time even if kinetics were twice as fast (*q* = 2). This is because the half-time of the receptors were 40 s (for *q* = 2), while the DA burst signal was repeated every 15 s. Thereby, [D1-DA] and [D2-DA] were dominated by the repetition of the signal rather than by the impact of individual DA burst signals. In contrast, for *q* =10 the change in receptor occupancy was dominated by the single pulses, since the half-life time was 8s, whereby the receptors mostly unbind in between DA pulses. Therefore, our results concerning the similarity of D1 and D2 receptors do not depend on the exact kinetics parameters or potential temperature effects, as long as the parameter changes are roughly similar for D1 and D2 receptors. However, DA receptor kinetics faster by a factor of 10 or more affected the ability of DA receptor occupancy to integrate DA pulses over time (**Fig. 9e, f**).

**Figure 9:**
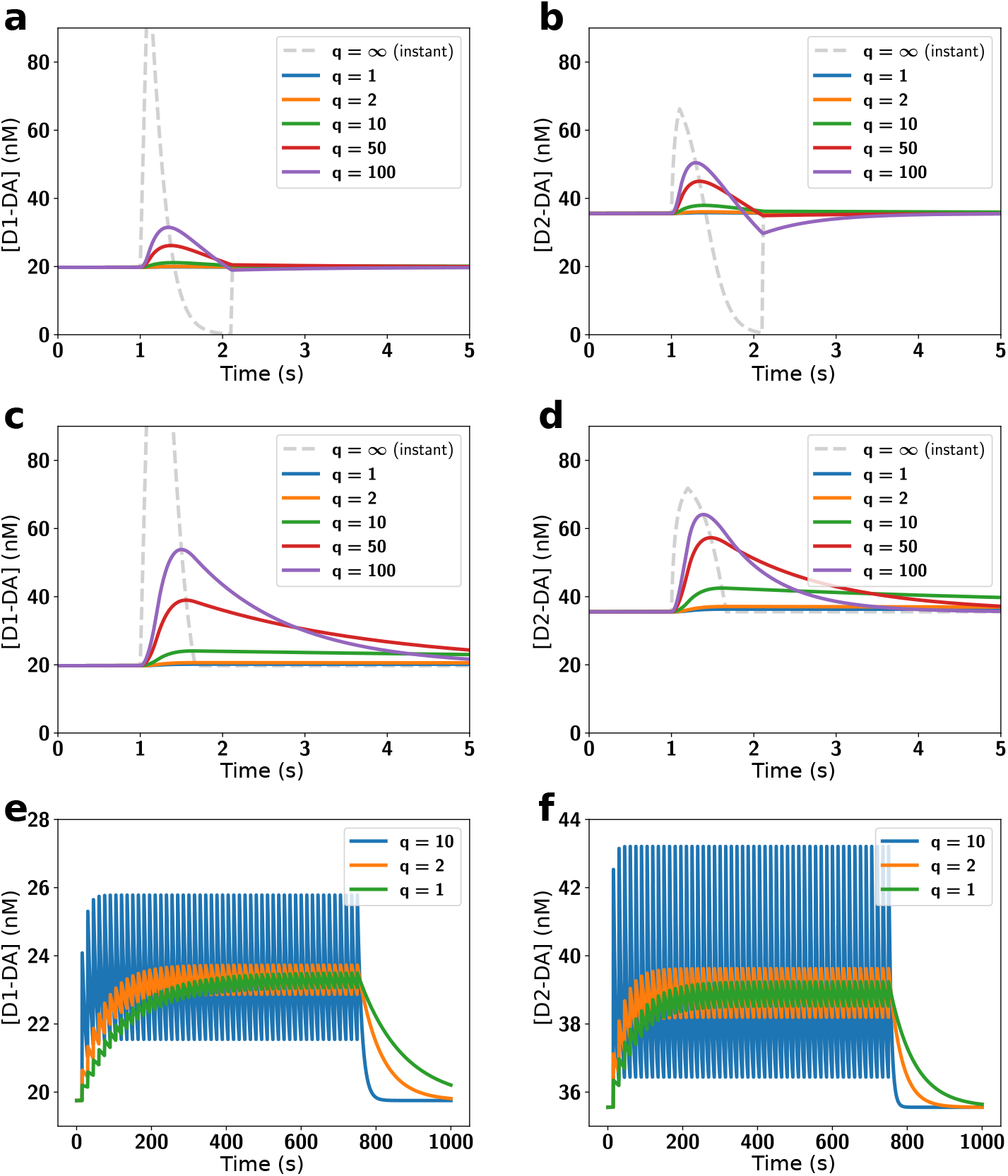
Similarities between D1 and D2 responses persist even if kinetics are much faster than our estimate. Absolute D1R occupancy ([D1-DA]; left column) and D2R occupancy ([D2-DA]; right column) were examined for burst-pause DA signals **(a, b)**, burst-only DA signals **(c, d)**, and the behavioural sequence **(e, f)** (i.e. same simulation scenarios as in Fig. 2a and the 50 bursts pattern from Fig. 4).

In our model we assumed homogeneous receptor populations, namely that all D1 receptors have a low affinity and that all D2 receptors have a high affinity. However, this could be a simplification, as ≈ 10% of D2 receptors may exist in a low affinity state, while ≈ 10% of the D1 receptors may be in a high affinity state (Richfield et al., 1989). Therefore, we also incorporated different affinity states of the D1 and D2 receptors into our model. The D1Rs in a high affinity state (*D*1^*high*^) were modelled by increasing the on-rate of the D1R but keeping its off-rate constant, creating a receptor identical to the *D*2^*high*^ receptor. Although the high affinity state kinetics of the D1R are currently unknown, we choose this model as a faster on-rate potentially has the strongest effect on our conclusions. Correspondingly, we modelled the *D*2^*low*^ receptor as a D2R with slower on-rate, which was equivalent to simply reducing [*D*2^*tot*^] since the *D*2^*low*^ receptors were predominantly unoccupied during baseline DA and bound only sluggishly to DA during phasic signals. The main effect of incorporating the different receptor affinity states was a change in the respective equilibrium values of absolute concentration of receptors bound to DA (**Fig. 10**). However, importantly, taking into account these different affinity states, preserved the similarity of time courses of D1R and D2R occupancy and the ability to integrate DA pulses over time (**Fig. 10** and **Fig. 11**) since the half-life time of both receptors remained long.

**Figure 10:**
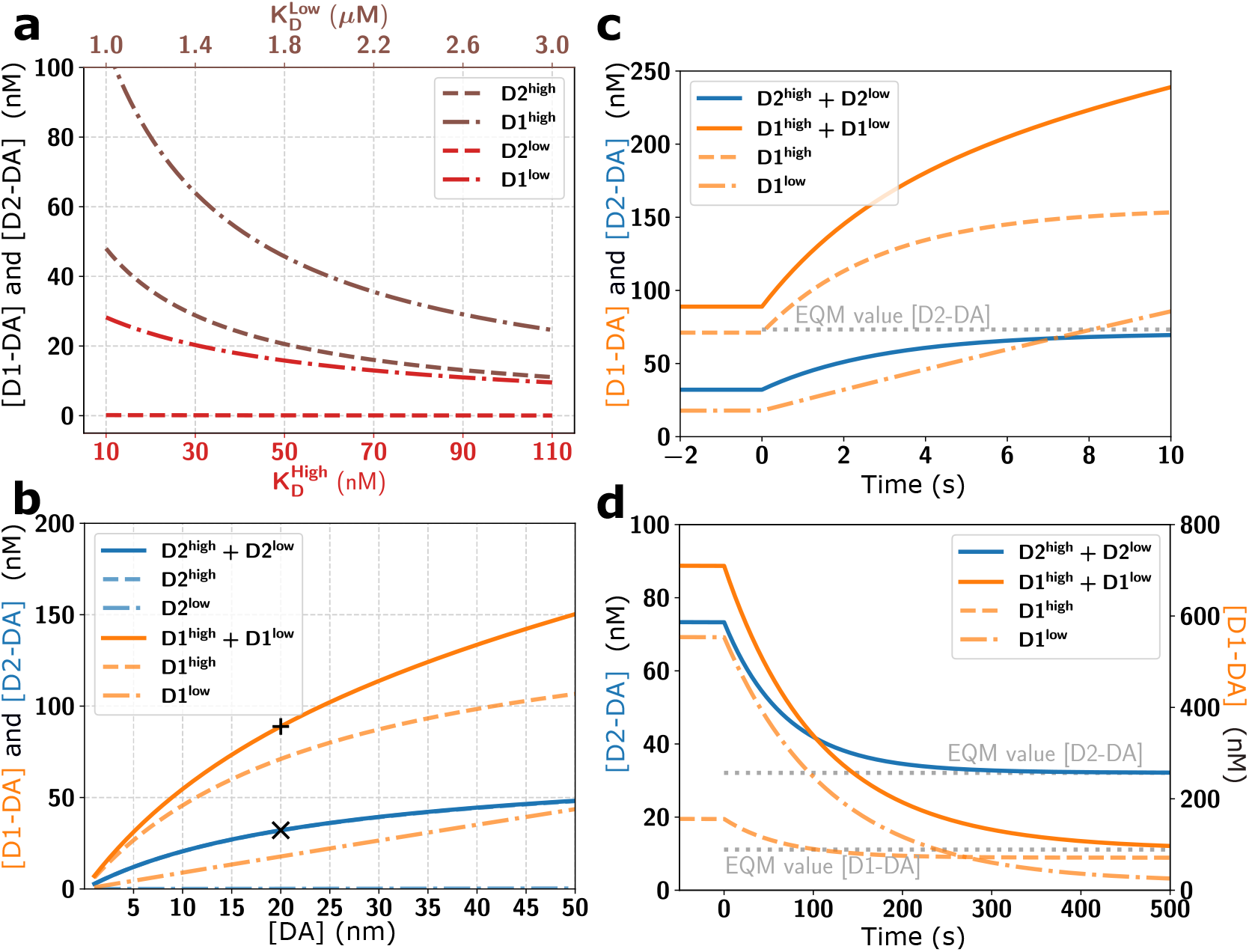
Baseline levels of D1 and D2 receptor occupation and impact of slow kinetics with different receptor affinity states. Here 10% of D1R are assumed to be in a high affinity state (*D*1^high^) and 90% of D1R in a low affinity state (*D*1^*low*^), while 10% of the D2R are in a low affinity state (*D*2^*low*^) and 90% of D2R are in their high affinity state (*D*2^*high*^). The overall receptor occupation for each receptor type is then the summed occupation of both states (*D*1^*high*^ + *D*1^*lo*w^ and *D*2^*high*^ + *D*2^*low*^). **(a)** The receptor occupancy at baseline [*DA*] = 20*nM* was dominated by the high affinity states for both receptors, even though only 10% of the D1R were in the high state. **(b)** The amount of bound D1R and D2R stayed within the same order of magnitude over a range of baseline [DA]. ‘×’ and +’ indicate the model default parameters. **(c)** As in the default model, for a large step up from [DA] = 20*nM* to [*DA*] = 1*μM*, and **(d)** a step down from [*DA*] = 1*μM* to [*DA*] = 20*nM*, D1 and D2 receptor occupancy approached their new equilibrium (EQM, grey dotted lines) only slowly (i.e. over seconds to minutes). As the [D1-DA] changes were dominated by the *D*1^*high*^ component, they were very similar to the D2R responses.

**Figure 11:**
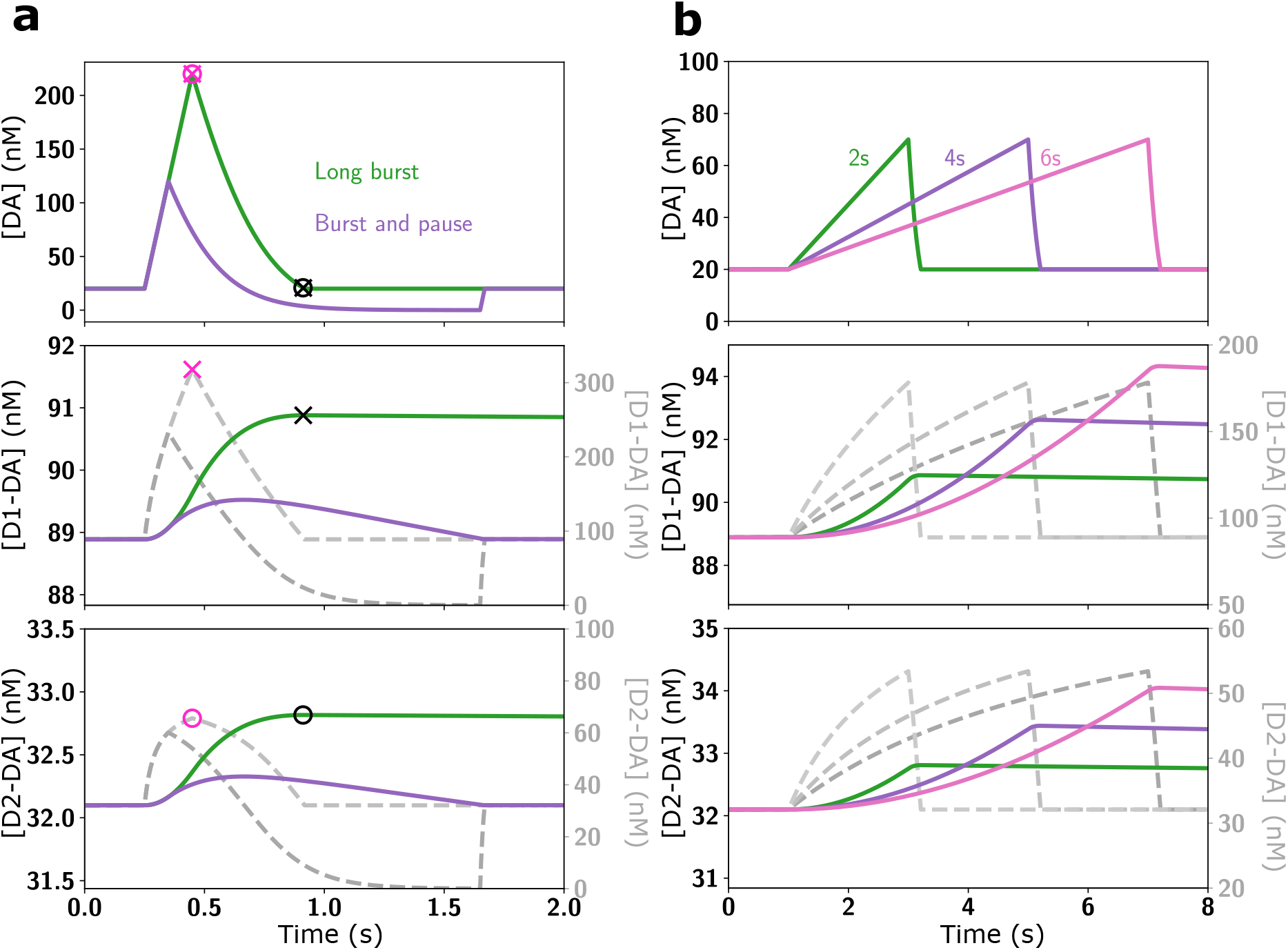
Impact of receptor kinetics on responses to different DA signals with 10% of D1R in a high affinity state (*D*1^*high*^) and 10% of D2 receptors in a low affinity state (*D*2^*low*^). **(a)** The effect of different phasic DA signals (top panels) on D1 (middle row) and D2 (bottom row) receptor occupancy in the slow kinetics model accounting for affinity states (coloured traces in middle and bottom panels; left scales) and to the affinity-based model (dashed grey traces, right scales). **(b)** Same as in the panel **a** but for DA ramps of different speed. As in the default model, the timing of the maximum receptor occupancy (‘×’ and ‘o’ for D1 and D2, respectively) coincides for instant kinetics (purple symbols) with the [DA] peak (combined x and o in top panel), while for slow kinetics (black symbols) it coincides with the offset of the [DA] signal instead (combined ‘×’ and ‘o’ in top row panel **a**). The main difference to the default model is the higher occupancy of the D1R, due to the *D*1^*high*^ component. There is no two-component unbinding since the *D*1^*high*^ and *D*1^*low*^ have similar off-rates, but differing on-rates. Overall, also for receptors with two affinity states, DA ramps are very effective in occupying the receptors.

## Discussion

The functional roles of DA in reward-related learning and motivation have typically been studied by characterizing the firing patterns of dopaminergic neurons and the resulting changes in striatal [DA] (Schultz, 2007). In contrast to other, more conventional neurotransmitters like glutamate or GABA, the release of DA in the striatum may form a global signal that affects large parts of the striatum similarly (Schultz, 1998). Such global [DA] changes involve longer time scales lasting at least several seconds (Roitman et al., 2008; Howe et al., 2013). Importantly, to affect neural activity in the striatum, DA first needs to bind to DA receptors. This process is often simplified by assuming that this happens instantaneously, so that every change in [DA] is immediately translated into a change in DA receptor occupation. As this contradicts physiological measurements of the receptor kinetics (Burt et al., 1976; Sano et al., 1979; Maeno, 1982; Nishikori et al., 1980), we developed and investigated a model incorporating DA receptor kinetics as well as differences in D1 and D2 receptor abundance in the striatum.

Our results cast doubt on several long-held views on DA signalling. A common view is that D1 and D2 MSNs in the striatum respond to different DA signals due to the affinity of their predominant receptor type. Accordingly, phasic DA changes should primarily affect D1 MSNs, while slower changes or DA pauses should primarily affect D2 MSNs. In contrast, our model indicates that both D1R and D2R systems can detect [DA] changes, independent of their timescale, equally well. That is, slow tonic changes in [DA], phasic responses to rewards, and ramping increases in [DA] over several seconds lead to a similar time course in the response of D1 and D2 receptor occupation in our model. However, the baseline level of activated DA receptors and the amplitude of the response was typically twice as high in D2 compared to D1 receptors. Although, D1 and D2 receptors have opposing effects on the excitability (Flores-Barrera et al., 2011) and strength of cortico-striatal synapses (Centonze et al., 2001), we challenge the view that differences in receptor affinity introduce additional asymmetries in D1 and D2 signalling. Instead of listening to different components of the DA signal, D1 and D2 MSNs may respond to the same DA input. This would actually increase the differential effect on firing rate responses of D1 and D2 MSNs because the opposite intracellular effects of D1 and D2 activation (Surmeier et al., 2007) occur then for the whole range of DA signals.

Recently, ramps in [DA], increasing over several seconds towards a goal, have been connected to a functional role of DA in motivation (Howe et al., 2013; Hamid et al., 2016). In our model DA ramps were very effective in occupying DA receptors due to their long duration. In contrast, for brief phasic increases, the receptor occupation was less pronounced. Overall, our model predicts that the area under the curve of DA signals determines the receptor activation, which puts more emphasis on the duration of the signals, rather than the amplitude of brief DA pulses.

Our model is also relevant for the interpretation of burst-pause firing patterns in DA neurons. These are a different firing pattern than the typical reward-related bursts, and consist of a brief burst followed by a brief pause in action potentials. Such two-component responses of DA cells may reflect saliency and value components, respectively (Schultz, 2016). For example, during extinction learning burst-pause firing patterns have been observed as a response to conditioned stimuli, with each component lasting about 100 ms (Pan et al., 2008). Our model provides a mechanistic account for how the burst-pause DA signals have a different effect on MSNs than pure burst signals, which is important to distinguish potential rewarding signals from other salient, or even aversive stimuli. In our model the pause following the burst was very effective in reducing the number of occupied receptors quickly, thereby preventing the otherwise long-lasting receptor occupation due to the burst. Thereby, canceling the effect of the brief burst might be a neural mechanism to correct a premature burst response that was entirely based on saliency rather than stimulus value (Schultz, 2016). As fast responses of DA cells to potentially rewarding stimuli are advantageous to quickly redirect behaviour, the subsequent pause signal might constitute an effective correction mechanism labelling the burst as a false alarm.

Functionally, the slow unbinding rate of D1 and D2 receptors pointed to an interesting property in integrating phasic DA events over time. The unbinding rate might be one of the mechanisms translating fast DA signals into a slower time scale, which could be a key mechanism to generate motivational signals (Mohebi et al., 2019). Importantly, the slow kinetics of receptor binding do not prevent a fast neuronal response to DA signals. In our model [DA] changes affected the number of occupied receptors immediately; it just took seconds or even minutes until the new equilibrium was reached. However, reaching the new equilibrium is not necessarily relevant on a behavioural level. Instead the intracellular mechanisms that react to the receptor activation need to be considered to determine which amount of receptor activation is required to affect neural activity. In our model changes on a nanomolar scale occurred within 100 ms, a similar timescale as behavioural effects of optogenetic DA manipulations (Hamid et al., 2016).

The slower time scales were introduced into our model by the kinetics based on in-vitro measurements (Burt et al., 1976; Sano et al., 1979; Maeno, 1982; Nishikori et al., 1980). A limitation of our model is the uncertainty about the accuracy of these measurements, and whether they reflect in-vivo conditions. We addressed this here by also examining faster kinetics, for which there is currently no direct evidence in the literature. However, recently DA receptors have been genetically modified to serve as sensors for fast [DA] changes (Patriarchi et al., 2018), which suggests possible fast kinetics. It seems unlikely though that the kinetics of the genetically-modified receptors represent the kinetics of the wildtype DA receptors, as e.g. the screening procedure to find suitable receptor variants yielded a large range of different affinities based on changes at the IL-3 site (Patriarchi et al., 2018). Changes in the IL-3 site have also previously been shown to strongly affect the receptor affinity (Robinson et al., 1994). Our broader view is that it is important to consider the effect of the receptor kinetics on DA signalling, which have not received much attention in experimental studies, nor in theoretical considerations of DA function so far.

## Acknowledgements

We thank Joshua Berke, Paul Overton, Alejandro Jimenez, Moham-madreza Mohagheghi Nejad and Amin Mirzaei for helpful discussions. This work was supported by the University of Sheffield and its high performance computing resources, by funding from the EU H2020 Programme as part of the Human Brain Project (HBP-SGA1, 720270; HBP-SGA2, 785907), and the BrainLinks-BrainTools Cluster of Excellence funded by the German Research Foundation (DFG, grant number EXC 1086), and the state of Baden-Wuerttemberg through bwHPC.

